# Within-host genotypic and phenotypic diversity of contemporaneous carbapenem-resistant *Klebsiella pneumoniae* from blood cultures of patients with bacteremia

**DOI:** 10.1101/2022.05.26.493675

**Authors:** Shaoji Cheng, Giuseppe Fleres, Liang Chen, Guojun Liu, Binghua Hao, Anthony Newbrough, Eileen Driscoll, Ryan K. Shields, Kevin M. Squires, Ting-yu Chu, Barry N. Kreiswirth, M. Hong Nguyen, Cornelius J. Clancy

## Abstract

Carbapenem-resistant *Klebsiella pneumoniae* (CRKP) are major pathogens globally. It is unknown whether bloodstream infections (BSIs) by CRKP and other bacteria are commonly caused by single organisms or mixed microbial populations. We hypothesized that contemporaneous CRKP from blood cultures of individual patients are genetically and phenotypically distinct. We determined short-read whole genome sequences of 10 strains from single colonies from CRKP-positive blood cultures in each of 6 patients (Illumina HiSeq). All strains were sequence type (ST)-258 *K. pneumoniae* that were unique by core genome single nucleotide polymorphism phylogeny, antibiotic resistance and virulence genes, capsular polysaccharide (CPS) gene mutations, and/or plasmid loss. Strains from each of 3 patients that differed in antibiotic resistance, virulence and/or CPS gene content underwent long-read sequencing for genome completion (Oxford Nanopore), and were tested for phenotypes *in vitro* and pathogenicity during mouse BSIs. Genetically distinct strains within individual patients exhibited significant differences in carbapenem, beta-lactam/beta-lactamase inhibitor and other antibiotic responses, CPS production, mucoviscosity, and susceptibility to serum killing. In 2 patients, strains differed significantly in their ability to infect organs and cause mortality in mice. In conclusion, we identified genotypic and phenotypic variant ST258 *K. pneumoniae* strains from blood cultures of individual patients, which were not detected by the clinical laboratory at time of BSI diagnosis. The data support a new paradigm of CRKP population diversity during BSIs. If validated for other BSIs, within-host bacterial diversity may have profound implications for medical, microbiology laboratory and infection prevention practices, and for understanding emergence of antibiotic resistance and pathogenesis.

**IMPORTANCE:** In processing positive microbiologic cultures, standard clinical laboratory practice is to test a single bacterial strain from each morphologically distinct colony. We performed comprehensive whole genome sequence analyses on 10 carbapenem-resistant *Klebsiella pneumoniae* (CRKP) strains from positive blood cultures from each of 6 patients. Our findings that all strains were genetically unique and that genetic variants manifested differences in phenotypes like antibiotic responsiveness and virulence suggest that CRKP bloodstream infections may be commonly caused by mixed bacterial populations. Results raise questions about laboratory protocols and treatment decisions that are directed against a single strain. The observation that pan-genome analyses revealed inter-strain differences that were not evident by studying core genomes has important implications for investigating nosocomial outbreaks and transmission. Data also suggest a model of pathogenesis of CRKP infections, in which environmental pressures *in vivo* may select for outgrowth of variants that manifest antibiotic resistance, tolerance or specific virulence attributes.

## INTRODUCTION

Carbapenem-resistant Enterobacterales (CRE) are “urgent threat” pathogens globally (1–3). *Klebsiella pneumoniae* are the most common CRE worldwide (4, 5). Most CRE infections are caused by commensal strains from the gastrointestinal (GI) tract (6–8). Recent whole genome sequence (WGS) data demonstrate that colonization or chronic infections by various bacteria may be caused by a population of clonal strains, in which genetic diversity emerges during long-term interactions with the host (9–22). Such clonal, but genetically diverse strains can manifest distinct phenotypes that are potentially relevant to enhanced commensalism or persistence in the host (9, 10). At present, it is unknown how commonly acute monomicrobial infections of normally sterile sites are caused by genetically and phenotypically diverse bacterial populations.

The prompt and accurate diagnosis of bacterial bloodstream infections (BSIs) is among the most critical functions of clinical microbiology laboratories (23). In processing positive microbiologic cultures, the standard practice in clinical laboratories is to isolate a single strain from each morphologically distinct colony. This practice is in keeping with the long-standing model of bacteremia, in which most monomicrobial infections are believed to be due to a single organism (“single organism” or “independent action” hypothesis) (24–29). Approximately 10-15% of BSIs are polymicrobial, with more than one species recovered from positive blood cultures (30). A smaller percentage of BSIs are monomicrobial but polyclonal, caused by strains of the same species that differ by multilocus sequence type (ST) or pulsed field gel electrophoresis patterns (31, 32). In this study, we tested the hypothesis that carbapenem-resistant *K. pneumoniae* (CRKP) from individual patients with putatively clonal, monomicrobial BSIs are genetically diverse and manifest phenotypic differences that are not recognized using routine clinical laboratory procedures. We identified 6 patients with CRKP BSIs, from whom the clinical laboratory isolated an index strain from a single colony morphotype. For each patient, we analyzed WGSs of the index strain and 9 other strains recovered from independent, morphologically indistinguishable colonies. We demonstrated that each strain from a given patient was a distinct genetic variant of ST258 *K. pneumoniae*, the predominant international clone. We assessed phenotypes of strains from 3 patents. We identified strains with distinct *in vitro* phenotypes in each patient, and strains with significant differences in virulence *in vivo* in 2 patients. Our findings support a new paradigm for understanding CRKP BSIs as diseases of diverse bacterial populations rather than single organisms.

## RESULTS

### WGSs of CRKP from positive blood cultures

We obtained first positive blood culture bottles from 6 adults who were diagnosed with monomicrobial CRKP BSI. The clinical microbiology laboratory identified a single colony morphotype in each case, from which a single index CRKP strain was isolated. We streaked aliquots from positive cultures onto blood agar plates, selected 9 colonies at random from each patient, and isolated a strain from each colony (Figure 1). We determined short-read WGSs of the index strain (labeled as strain 1) and other strains (labeled as strains 2-10) for each patient (Illumina HiSeq). All strains were identified as ST258 *K. pneumoniae*. Information on genome assemblies is provided in Supplemental Table 1. *K. pneumoniae* 30660/NJST258_1 served as reference genome for WGS analyses.

**Figure 1.**
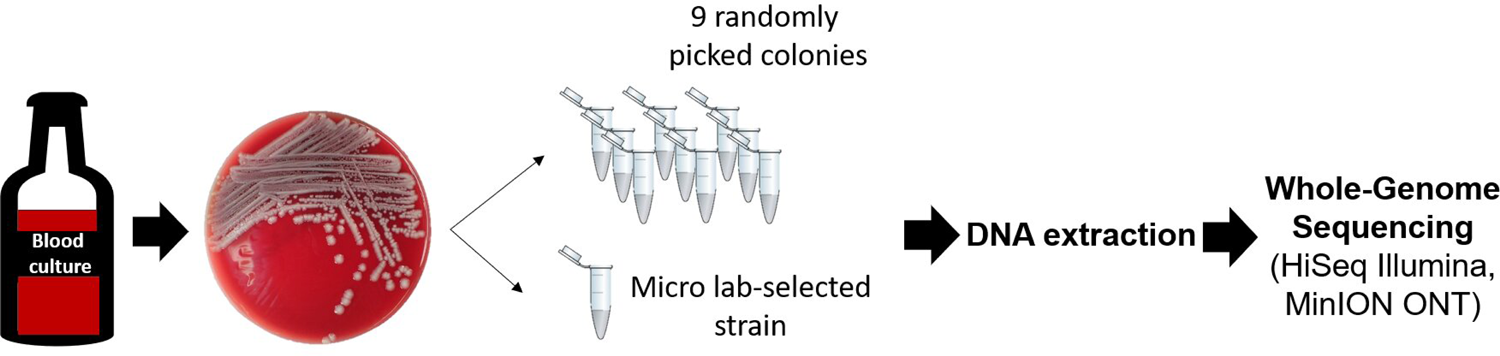
Selection of carbapenem-resistant *Klebsiella pneumoniae* strains. First positive blood culture bottles from each of 6 patients diagnosed with carbapenem-resistant *Klebsiella pneumoniae* bloodstream infection were obtained from the University of Pittsburgh Medical Center clinical microbiology laboratory immediately after routine microbiological work-up. An aliqout from culture bottles was streaked on blood agar plates, which were cultured overnight. For every patient, we isolated and randomly selected a single strain from 9 independent colonies. In addition, we collected the strain isolated by the clinical laboratory (index strain). DNA extracted from these 10 strains per patient underwent whole genome sequencing (Illumia HiSeq). Frozen stocks were made of each colony and pooled growth from each plate. WGS: Whole genome sequencing

### Core genome single nucleotide polymorphism (SNP) phylogeny

To estimate genetic relationships among strains, we first performed core genome SNP analysis. An alignment of core genome nucleotides was used to build a high-resolution SNP phylogenetic tree. Strains segregated into clade 1 (capsule type KL106; patients B and G) and clade 2 (capsule type KL107; patients A, D, F and J) (Figure 2A). Each patient’s strains clustered closely on the phylogenetic tree. Clade 1 strains differed by 0-1173 SNPs; inter-patient differences for strains from patients B and G were ≥144 SNPs (Figure 2B). Clade 2 strains differed by 0-50 SNPs; inter-patient differences for strains within clade 2 were ≥17 SNPs. Within-host differences between several strains from patients A and G were ≥15 SNPs, a cut-off for *K. pneumoniae* strain identity determined in genomic epidemiology studies at our center and others (33).

**Figure 2.**
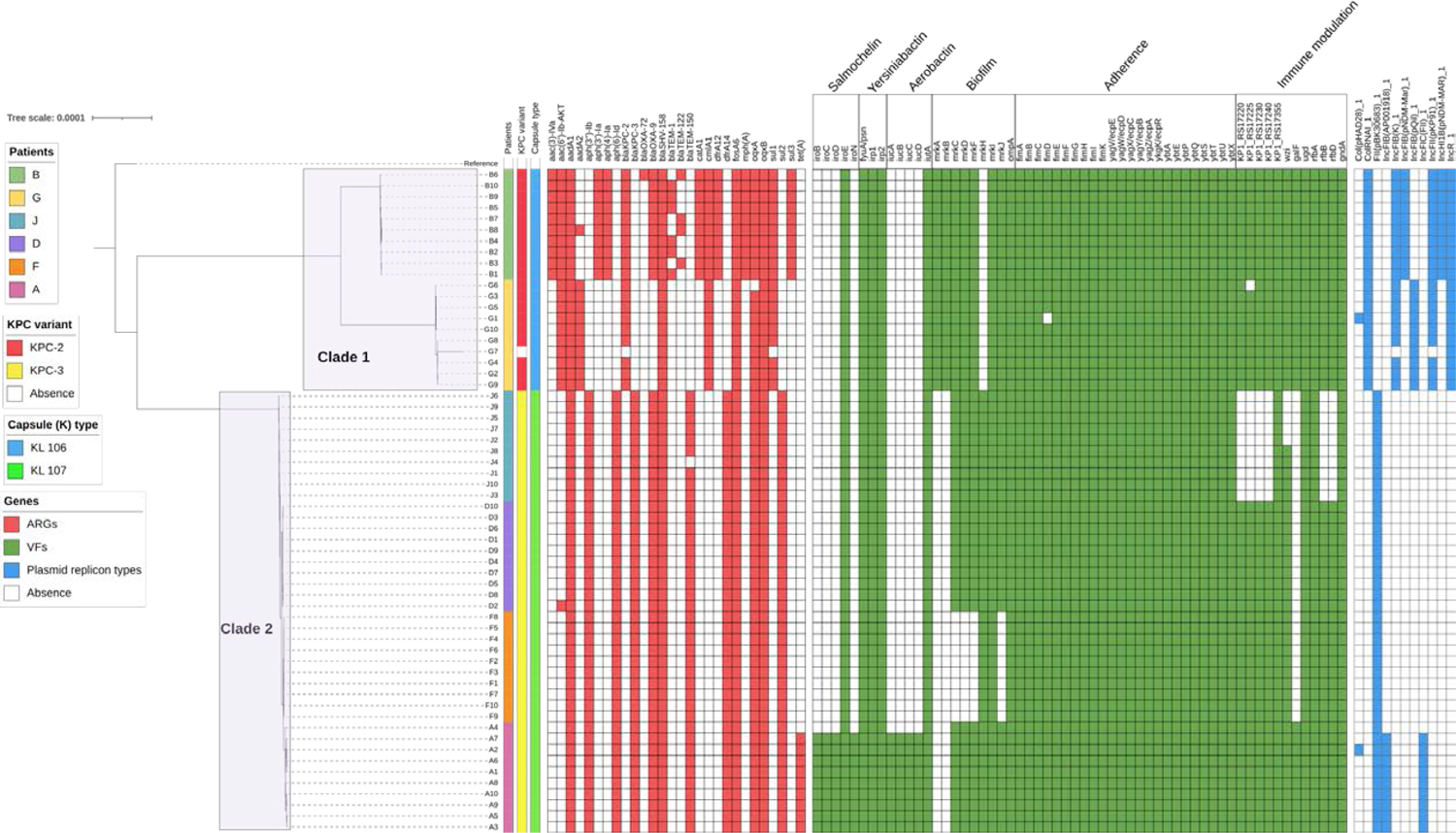
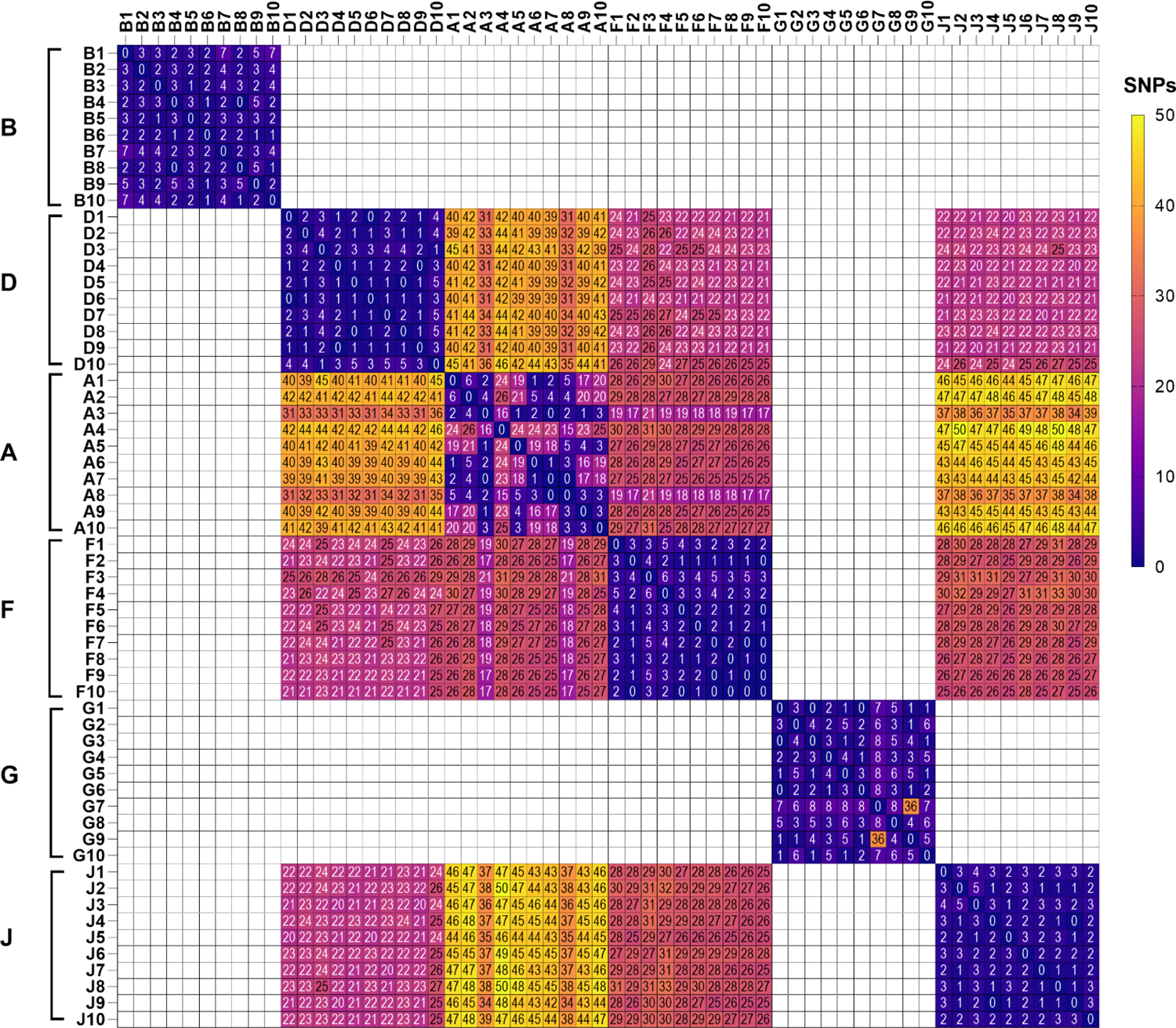
Comparisons of carbapenem-resistant *Klebsiella pneumoniae* strains by core genome single nucleotide polymorphism (SNP) phylogeny, and presence/absence of specific genes. **Figure 2A. Summary of strain comparisons.** Data were generated by analyses of short-read whole genome sequences (Illumina HiSeq). The figure legend appears on the left. From left-to-right, the following comparisons between strains are presented: core genome SNP phylogenetic tree, carbapenemases, capsule (KL) types, and presence/absence of specific antibiotic resistance (red), virulence (green) and plasmid replication type (blue) genes. The reference strain *was K. pneumoniae* 30660/NJST258_1 (GenBank assembly accession: GCA_000598005.1). Taken together, these comparisons demonstrated within-host genetic diversity among strains from 5 of 6 patients (A, B, D, G, J; no differences demonstrated for patient F. **Figure 2B. Heat map of core genome SNPs.** Brackets on the left of the figure demark strains from the respective patients. Numbers of core genome SNP differences are within matrix boxes. Within-host differences among several strains from patients A and G were ≥ 15 core genome SNPs, a cut-off for strain identity defined in earlier studies at our center (33).

### Antibiotic resistance, capsular biosynthesis, virulence and plasmid replicon genes

We next compared specific genome content of strains by surveying short-read WGS data for presence of genes involved in antibiotic resistance, capsular biosynthesis and virulence, and for genes associated with plasmid replicon types (Figure 2A). Within-host diversity in resistance gene content was evident among strains from 5 patients (A, B, D, G, J). All strains except G7 carried *bla*_KPC_. Clade 1 and 2 strains carried *bla*_KPC-2_ and *bla*_KPC-3_, respectively.

Within-host diversity in capsular or other virulence gene content was evident in strains from 5 patients (A, D, F, G, J) (Table 1, Figure 2A). In 3 of these patients (A, D, J), within-host diversity was evident in capsular genes. All 10 strains in patients D, F and J were missing *galF*, which encodes an alpha-D-glucosyl-1-phosphate uridylyltransferase involved in translocation and surface assembly of capsule (34). Strain D10 was also missing capsular genes *wzi, wza and wzb*. Strains from patient J had deletions of various capsular genes (discussed in detail below). At least one strain from patients A and J carried substitutions or frame-shift mutations in *wzc*.

**Table 1.**
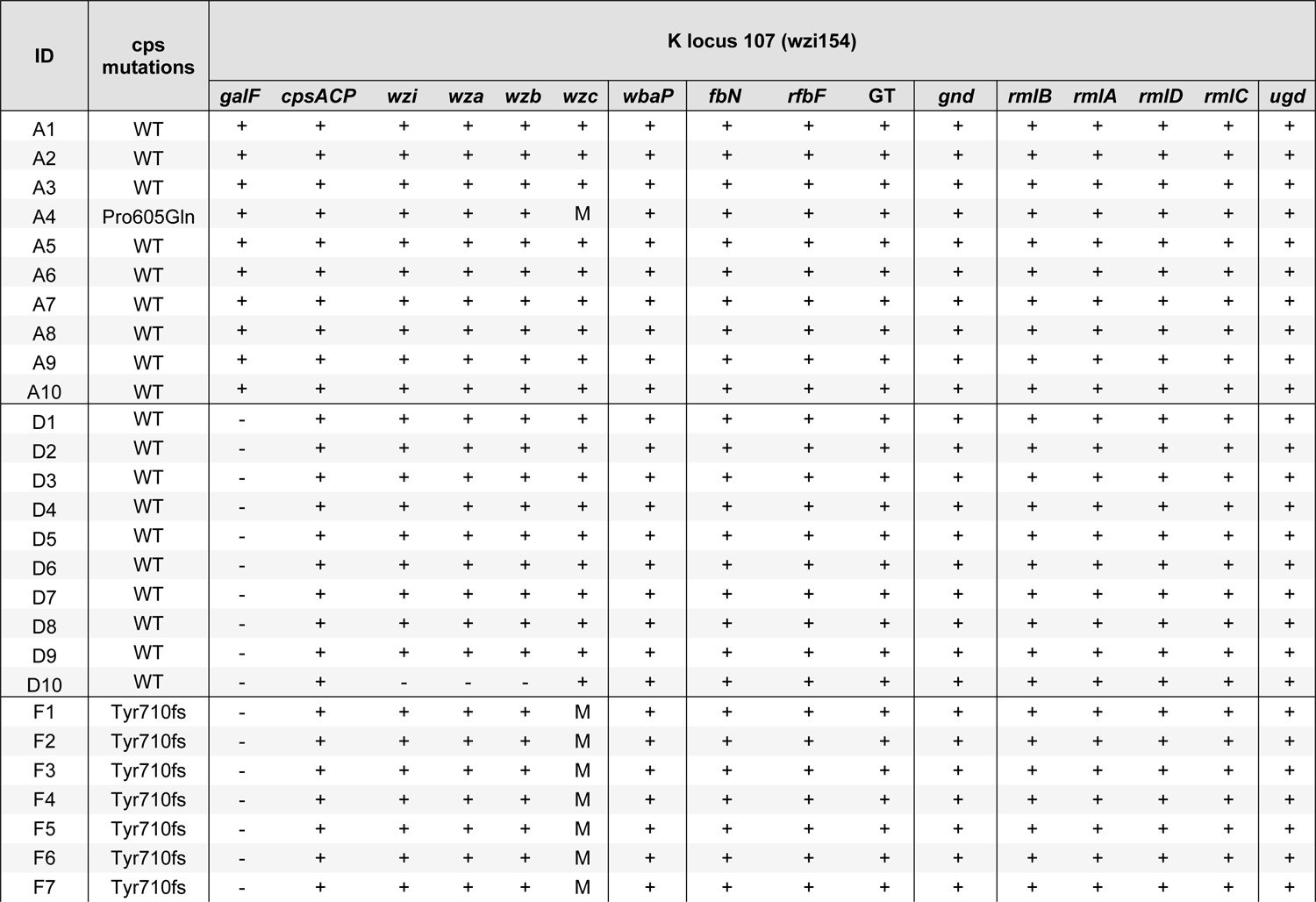
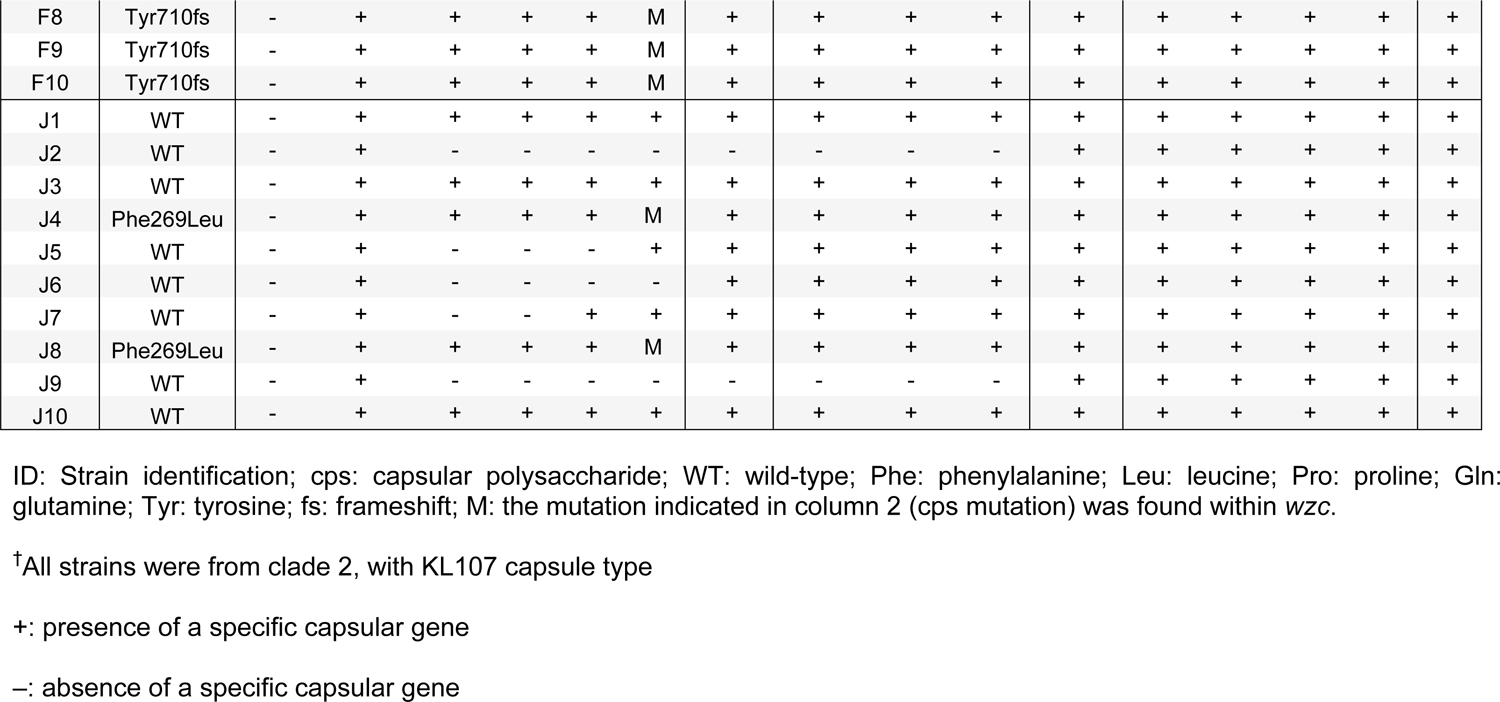
Carbapenem-resistant *Klebsiella pneumoniae* strains with capsular gene mutations.

Within-host diversity in plasmid replicon types was evident among strains from 2 patients (A, G). Clade 1 and 2 strains had genes associated with 8 (IncF, n=4; Col, n=2; IncH, n=1; IncR, n=1) and 4 plasmid replicon types (IncF, n=3; Col, n=1), respectively.

### Pan-genome analyses

To complete short-read WGS comparisons, we performed pan-genome analyses by constructing presence/absence matrices for 7062 genes, including 4700 core genes (present in ≥60 of 61 genomes), 201 soft-core genes (57-59 genomes) and 2161 accessory genes (1-56 genomes) (Figure 3; Supplemental Table 2). Strains segregated by clade and by patient in the accessory gene phylogenetic tree. In each patient, within-host diversity was detected among strains, including differences that were not evident by preceding short-read WGS analyses.

**Figure 3.**
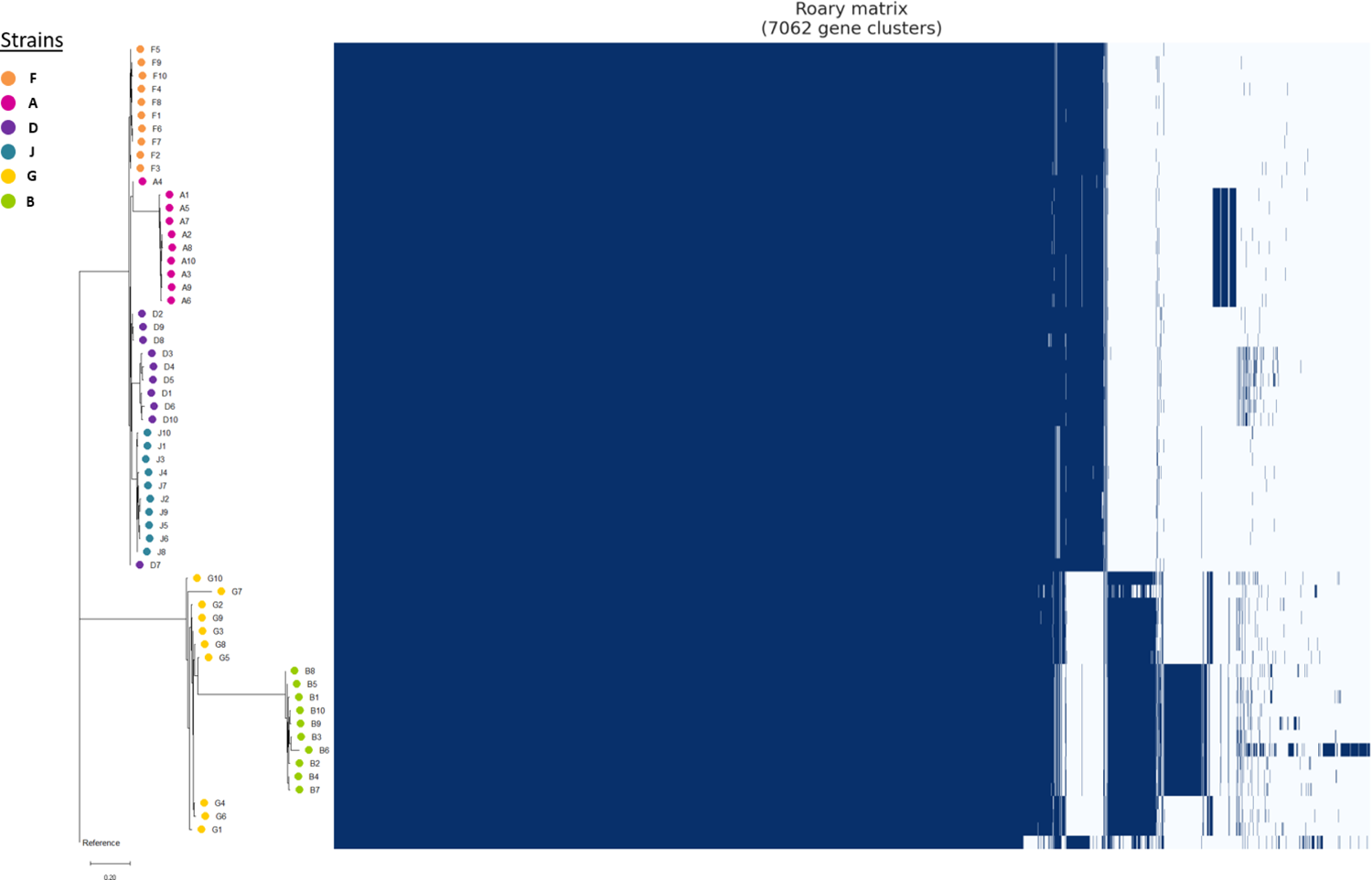
Pan-genome comparisons of carbapenem-resistant *Klebsiella pneumoniae* strains. Data were generated by analyses of short-read whole genome sequences (Illumina HiSeq). The phylogenetic tree was constructed using data on presence/absence of 7062 gene clusters, including 4700 core genes (present in 60-61 genomes), 201 soft-core genes (present in 57-59 genomes) and 2161 accessory genes (present in 1-57 genomes). The gene presence/absence matrix is shown to the right of the phylogenetic tree. Blue: gene presence; white: gene absence. Within-host genetic diversity of each strain in all 6 patients was evident by pan-genome analyses. Please refer to Supplementary Table 2 (Excel file) for sequence data for gene presence/absence matrices.

### Detailed descriptions of within-host CRKP genetic diversity in three patients

In the remainder of the study, we investigated strains from patients A, G and J. We focused on these patients since strains showed within-host differences in antibiotic resistance and virulence genes.

#### Patient A

Strain A4 was the most distinct A strain by short-read WGS analyses. A4 differed from other A strains by an average of 22 core genome SNPs. It was unique among A strains for lack of *tetA* (tetracycline resistance), virulence genes encoding for aerobactin (*iucABCD*) and the salmochelin siderophore system (*iroBCDN*), and plasmid replicons IncFIB and IncFIC (Figures 2A, 2B).

We performed long-read sequencing (Oxford Nanopore WGS, MinION) on strains A1 and A4, and generated hybrid assemblies with short-read WGS data to obtain complete chromosome and plasmid sequences. A1 and A4 were chosen since they were the index strain and most distinct strain by short-read analyses, respectively. We identified SNPs or insertions/deletions (indels) in 25 genes, including 16 and 9 that resulted in non-synonymous and synonymous differences, respectively; a SNP and an indel were also identified in intergenic regions (Table 2). Notable non-synonymous mutations were observed in *wzc* (P605Q substitution), porin *ompK36* (586 C>T; pre-mature stop codon), ferric iron reductase *fhuF* (G28A), and fimbrin adhesin *fimH* (G287A). Long-read data confirmed that A4 lacked a 160 kb IncFI plasmid that carried *tetA* and several virulence genes (Table 3).

**Table 2.**
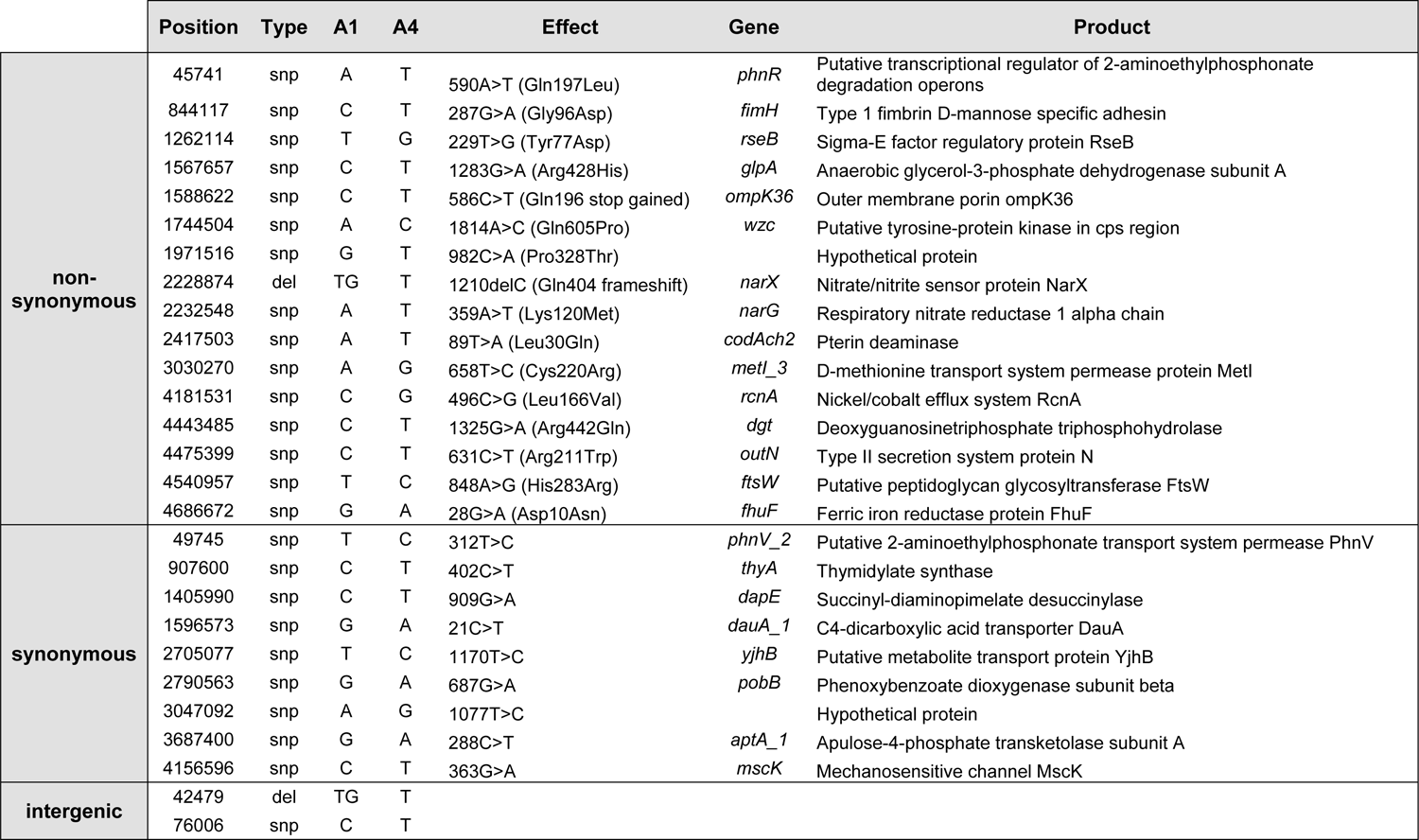
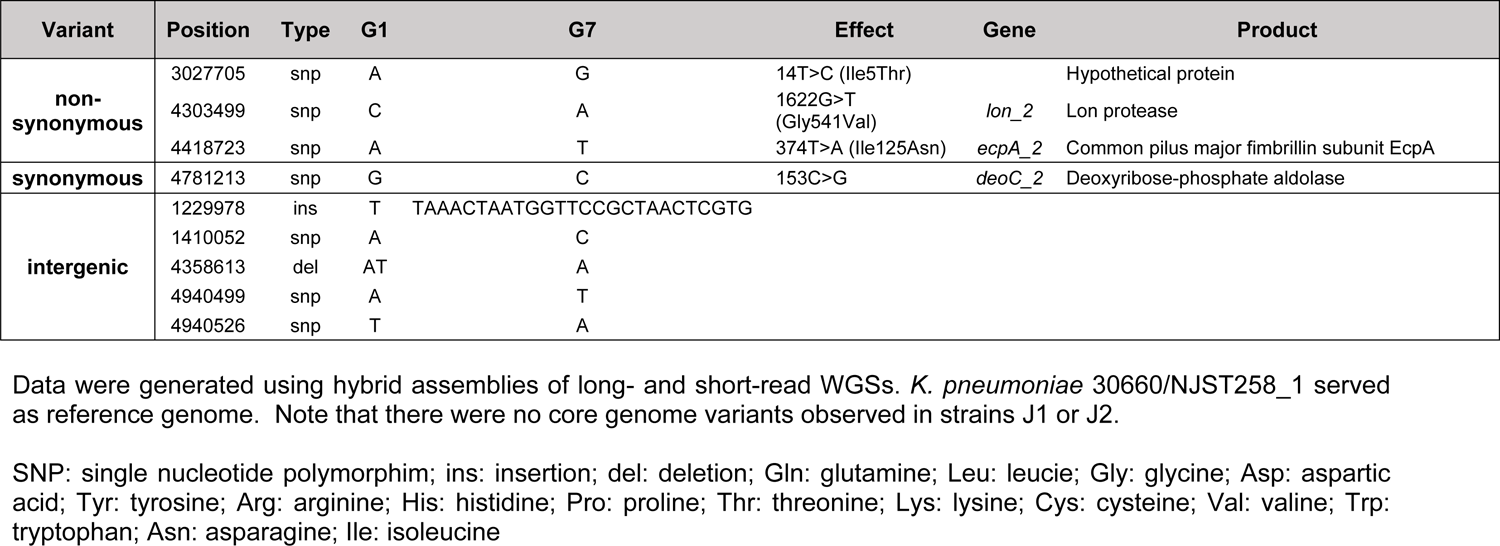
Core genome single nucleotide polymorphisms and insertion-deletions in carbapenem-resistant *Klebsiella pneumoniae* strains from patients A (A1 and A4) and G (G1 and G4).

**Table 3.** Pan-genome comparisons of carbapenem-resistant *Klebsiella pneumoniae* from three patients (A G, J).

| Strain | Contig | Length (bp) | Plasmid Replicons | Resistance genes | Virulence genes |
| --- | --- | --- | --- | --- | --- |
| A1 | 1 (Chr) | 5,499,994 |  | *blaSHV-158, fosA6, oqxA, oqxB* | *acrAB, ecpABCDER, entABCEFS, fepABCDG, fimABCDEFGHGIK, mrkCDFJIH, rpoS, rcsB, galF, gndA, ugd, rfbABD, clbABCDEFGHGIJKLMNOPQS, fyuA/psn, irp1, irp2, rcsA, iroE, ybtAETUPQXS, tssBCDFGHJKLM, iutA, ompA, fur, fes, wzi, KP1_RS17220, KP1_RS17225, KP1_RS17230, KP1_RS17240, KP1_RS17355* |
|  | 2 (Plasmid) | 166,337 | IncFIC(FII), IncFIB(AP001918) | *tet(A)* | *iroBCDEN, iutA, iucABCD* |
|  | 3 (Plasmid) | 161,666 | FII(pBK30683) | *blaKPC-3, blaTEM-150, blaOXA-9, blaSHV-158, dfrA14, sul2, aph(3'')-Ib, aph(6)-Id, aadA1, aac(6')-Ib-AKT* |  |
| A4 | 1 (Chr) | 5,498,521 |  | *blaSHV-158, fosA6, oqxA, oqxB* | *acrAB, ecpABCDER, entABCEFS, fepABCDG, fimABCEFGHIK, mrkCDFJIH, rpoS, rcsB, galF, gndA, ugd, rfbABD, clbABCDEFGHGIJKLMNOPQS, fyuA/psn, irp1, irp2, rcsA, iroE, ybtAETUPQXS, tssBCDFGHJKLM, iutA, ompA, fur, fes, wzi, KP1_RS17220, KP1_RS17225, KP1_RS17230, KP1_RS17240, KP1_RS17355* |
|  | 2 (Plasmid) | 161,665 | FII(pBK30683) | *blaKPC-3, blaTEM-150, blaOXA-9, blaSHV-158, dfrA14, sul2, aph(3'')-Ib, aph(6)-Id, aadA1, aac(6')-Ib-AKT* |  |
| G1 | 1 (Chr) | 5,550,531 |  | *blaSHV-158, fosA6, oqxA, oqxB, sul1, aadA2* | *acrAB, ecpABCDER, entABCEFS, fepABCDG, fimABCDEFGHGIK, mrkABCDFJI, rpoS, rcsB, galF, gndA, ugd, rfbABD, clbABCDEFGHGIJKLMNOPQS, fyuA/psn, irp1, irp2, rcsA, iroE, ybtAETUPQXS, tssBCDFGHJKLM, iutA, ompA, fur, fes, wzi, KP1_RS17220, KP1_RS17225, KP1_RS17230, KP1_RS17240, KP1_RS17355* |
|  | 2 (Plasmid) | 167,851 | IncFII(pKP91), IncFIB(K) | *blaKPC-2* |  |
|  | 3 (Plasmid) | 126,321 | IncR, IncFIB(pQil) | *aac(6')-Ib-AKT, aadA1, aadA2, cmlA1* |  |
|  | 4 (Plasmid) | 9,294 | ColRNAI |  |  |
| G7 | 1 (Chr) | 5,542,117 |  | *blaSHV-158, fosA6, oqxA, oqxB* | *acrAB, ecpABCDER, entABCEFS, fepABCDG, fimABCDEFGHGIK, mrkABCDFJI, rpoS, rcsB, galF, gndA, ugd, rfbABD, clbABCDEFGHGIJKLMNOPQS, fyuA/psn, irp1, irp2, rcsA, iroE, ybtAETUPQXS, tssBCDFGHJKLM, iutA, ompA, fur, fes, wzi, KP1_RS17220, KP1_RS17225, KP1_RS17230, KP1_RS17240, KP1_RS17355* |
|  | 2 (Plasmid) | 126,542 | IncR, IncFIB(pQil) | *aac(6')-Ib-AKT, aadA1, aadA2, cmlA1* |  |
|  | 3 (Plasmid) | 9,294 | ColRNAI |  |  |
| **J1** | 1 (Chr) | 5,389,279 |  | *blaSHV-158, fosA6, oqxA, oqxB* | *acrAB, ecpABCDER, entABCEFS, fepABCDG, fimABCDEFGHJK, mrkCDFJIH, rpoS, rcsB, gndA, ugd, rfbA, clbABCDEFGHJKLMNOPQS, fyuA/psn, irp1, irp2, rcsA, iroE, ybtAETUPQXS, tssBCDFGHIJKLM, iutA, ompA, fur, fes, wzi, KP1_RS17355* |
|  | 2 (Plasmid) | 165,212 | FII(pBK30683) | *blaKPC-3, blaTEM-150, blaOXA-9, blaSHV-158, dfrA14, sul2, aph(3'')-Ib, aph(6)-Id, aadA1, aac(6')-Ib-AKT* |  |
|  | 3 (Chr) | 93,071 |  |  |  |
| **J2** | 1 (Chr) | 5,377,661 |  | *blaSHV-158, fosA6, oqxA, oqxB* | *acrAB, ecpABCDER, entABCEFS, fepABCDG, fimABCDEFGHJK, mrkCDFJIH, rpoS, rcsB, gndA, ugd, rfbA, clbABCDEFGHJKLMNOPQS, fyuA/psn, irp1, irp2, rcsA, iroE, ybtAETUPQXS, tssBCDFGHIJKLM, iutA, ompA, fur, fes, KP1_RS17355* |
|  | 2 (Plasmid) | 165,212 | FII(pBK30683) | *blaKPC-3, blaTEM-150, blaOXA-9, blaSHV-158, dfrA14, sul2, aph(3'')-Ib, aph(6)-Id, aadA1, aac(6')-Ib-AKT* |  |
|  | 3 (Chr) | 93,071 |  |  |  |
|  | **4 (Plasmid)** | **4,097** |  |  |  |

Data were generated using hybrid assemblies of long- and short-read WGSs. Plasmid information was mined using long-read WGS. Within-host differences in content are highlighted as bolded text.
Chr: chromosome; bp: basepairs
KP1\_RS17220: glycosyltransferase, LPS; KP1\_RS17225: glycosyltransferase family 4 protein, LPS; KP1\_RS17230: glycosyltransferase, LPS; KP1\_RS17240: DUF4422 domain-containing protein, LPS; KP1\_RS17355: phosphatase PAP2 family protein, capsule

#### Patient G

Strain G7 was the most distinct G strain by short-read WGS analyses. G7 differed from other G strains by an average of 10 core genome SNPs, and in its lack of *bla_KPC-2_* and *sul1* (sulfonamide) resistance genes (Figure 2A). Strains G4 and G6 were notable for an *IS*5 insertion in the promotor region of *ompK36*, which has been shown to impact some broad-spectrum beta-lactam minimum inhibitory concentrations (MICs) (Supplemental Figure 1) (35). Strains G1 and G6 lacked *fimD* (encoding an adhesin) and gene KP1_RS17225 (encoding for a glycosyltransferase family 4 protein in the CPS synthesis region), respectively (Figure 2A); otherwise, virulence gene content was similar among G strains. Strain G7 lacked IncFIB and IncFII plasmid replicons.

Long-read sequencing was performed and hybrid assemblies were constructed for strains G1 (index strain) and G7 (most distinct strain by short-read WGS analyses). We identified SNPs in 4 genes, 3 of which were non-synonymous (Table 2); 5 other mutations were identified in intergenic regions. Strain G7 was confirmed to lack a 167,851 bp IncF1K plasmid with IncFIB and IncFII replicons that carried *bla*_KPC-2_ (Table 3).

#### Patient J

Strains from patient J were closely related by core-genome SNP analysis, with differences of 0-5 SNPs. Strain J4 was unique in lacking *bla*_TEM-150_, which encodes an extended spectrum beta-lactamase. J strains manifested 6 variations of KL107 capsule, associated with deletions and mutations of capsular biosynthesis genes (*galF*, *wzi*, *wbc*, *wzc*, *wbaP*) (Table 1; Figure 4). The predominant capsule had a 2.2 kb deletion encompassing *galF* (ΔKL 107-2.2kb; J1, J3, J10). Strains J4 and J8 had this 2.2 kb deletion, as well as Phe269Leu substation in *wzc*. Additional capsular gene deletions were detected in other J strains: ΔKL 107-2.2kb+2.7kb (J7), ΔKL 107-2.2kb+3.6kb (J5), ΔKL 107-2.2kb+6kb (J6), and ΔKL 107-2.2kb+11kb (J2, J9). The same IncFII (pBK30683) plasmid replicon was shared among all J strains (36).

**Figure 4.**
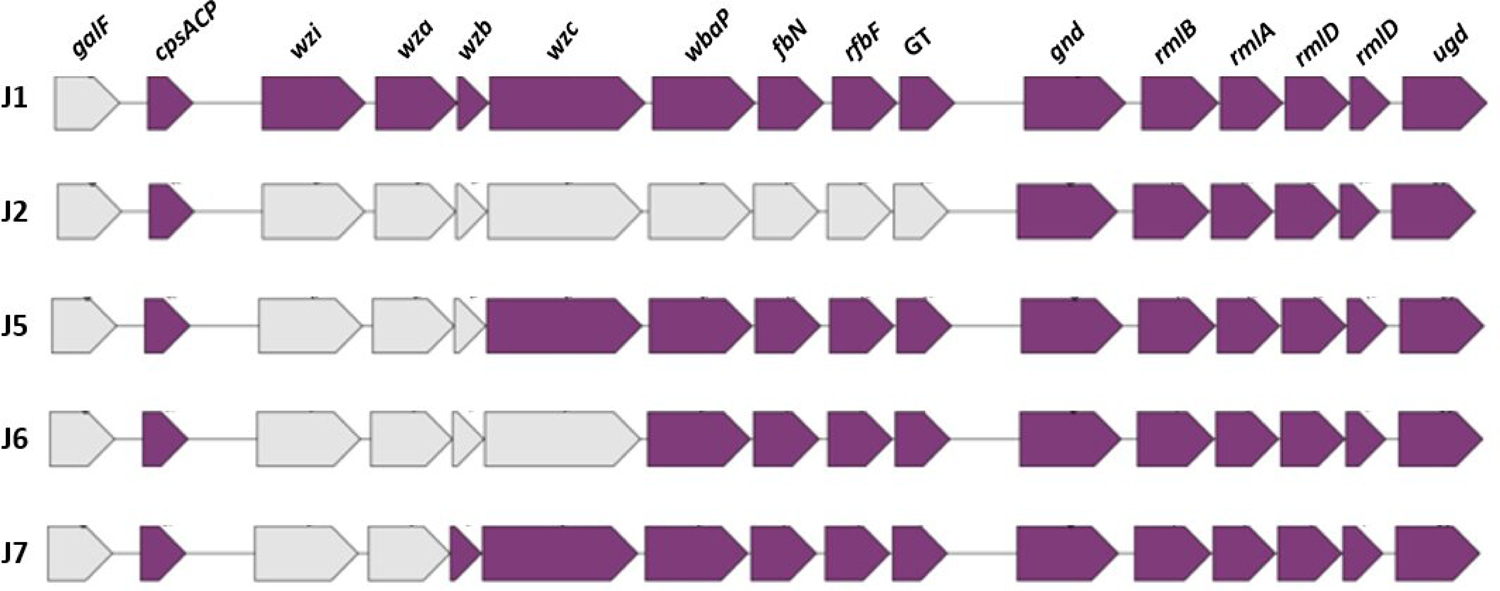
Capsular gene composition of carbapenem-resistant *Klebsiella pneumoniae* strains from patient J. Data are shown for representative J strains with given capsular gene composition. Based on gene deletions, J strains fell into 5 groups: 1) J1, J3, J4, J8, J10; 2) J2, J9; 3) J5; 4) J6; 5) J7. In addition to deletions shown in the figure, strains J4 and J8 also carried Phe262Leu substitutions in *wzc*. Therefore, J strains manifested 6 variations of KL107 capsule Grey and purple shading indicate gene absence and presence, respectively. Please refer to Table 1 for further details on capsular gene mutations in strains from patients A, D, F and J.

We chose J1 and J2 for long-read sequencing since they were the index strain and strain with the largest capsular deletion, respectively. Hybrid assemblies did not reveal SNPs or indels. Strain J2 carried a unique 4097 bp sequence that did not include known antibiotic resistance or virulence genes. Best match with this sequence by Blast nucleotide search was *E. coli* OK15 plasmid unnamed4 (12273 bp; GenBank: CP081681.1; 100% query coverage; 83.62% identity).

### Phenotypes of CRKP from three patients

There were no significant within-host differences in growth rates of A, G or J strains at either 30°or 37°C in Mueller-Hinton medium or M9 minimal medium without or with 100 µM iron chelator deferoxamine (data not shown).

#### Antibiotic susceptibility

Antibiotic MICs against A, G and J strains are presented in Supplemental Table 3. Strain A4 was susceptible (MIC: 4 μg/mL) to tetracycline whereas other strains were resistant (MIC: 256 μg/mL), consistent with absence and presence of *tetA*, respectively. Strain G7 was unique among G strains for susceptibility to meropenem (consistent with absence of *bla*_KPC-2_) and ceftazidime. Strains G4 and G6, which had an *IS*5 insertion in the promotor region of *ompK36*, exhibited meropenem-vaborbactam MICs that were ≥4-fold higher than those exhibited by other G stains. *ompK36* expression by G4 and G6 was reduced ∼60-fold and ∼100-fold (*p*<0.0001 for either strain), respectively, compared to other G strains by reverse transcription-polymerase chain reaction in both presence and absence of meropenem-vaborbactam; ompK36 production was significantly diminished in strains G4 or G6, as evident by sodium dodecyl sulfate–polyacrylamide gel electrophoresis (Supplemental Figure 2).

For J strains, there were no significant differences in MICs (Supplemental Table 3). In a recent study, a capsule deficient *K. pneumoniae* mutant strain had greater survival than a wild-type strain within bladder epithelial cells *in vitro* following exposure to meropenem-vaborbactam, in absence of phenotypic resistance (37). We performed 24 hour meropenem-vaborbactam and ceftazidime-avibactam time-kill experiments against strains J1 and J2, which were susceptible to the drugs by MICs (Figure 5). These drugs are the currently preferred treatment options against CRKP and other CRE infections (38). There was no difference between strains in initial kills at 4 hours by either agent. However, between 4-24 hours post-exposure, J2 regrew in presence of meropenem-vaborbactam and ceftazidime-avibactam; in contrast, there was no regrowth of J1 (39). At 24 hours, J2 concentrations were significantly greater than those of J1 (mean log_10_ CFU ± standard error: 9.9±1.8 *vs.* 1.8±0.3, respectively, for meropenem-vaborbactam at 4x MIC, p=0.03; 7.5±0.04 *vs.* 0.32±0.32 for ceftazidime-avibactam at 1x MIC, p=0.03). J2 isolates recovered after 24 hours did not demonstrate increases in meropenem-vaborbactam or ceftazidime-avibactam MICs.

**Figure 5.**
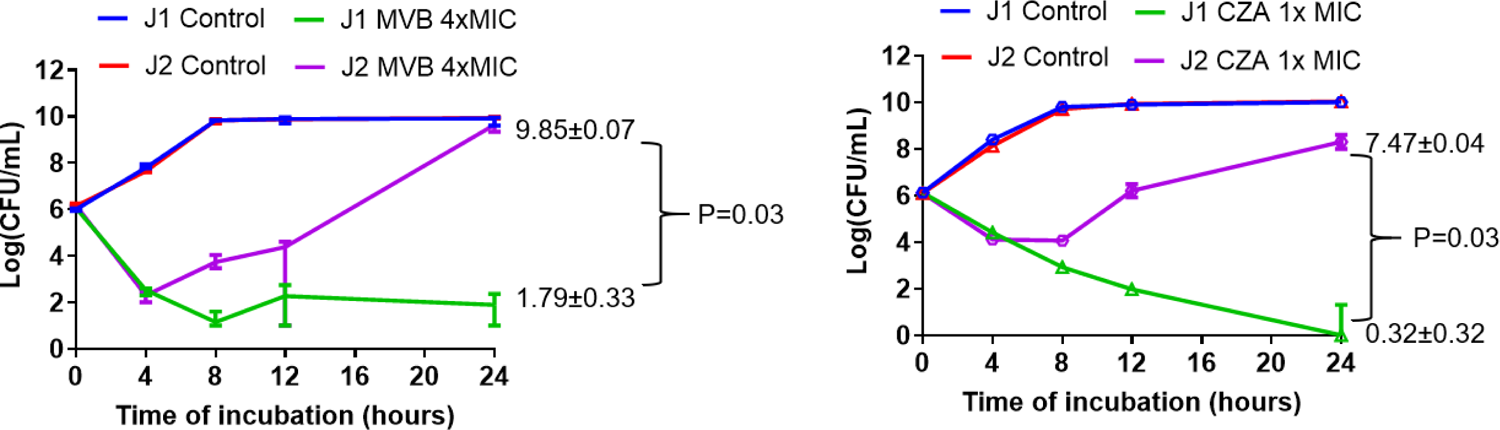
Meropenem-vaborbactam (MVB) and ceftazidime-avibactam (CZA) time-kills of carbapenem-resistant *KIebsiella pneumoniae* strains J1 and J2. MVB time-kills of strains J1 and J2 at 0x and 4x MIC are shown on the left. CZA time-kills at 0x and 1x MIC are shown on the right. Strains were incubated with 1x and 4xMIC of both drugs. For clarity in figures, data are not shown for MVB at 1x MIC or CZA at 4x MIC. MVB time-kills of J1 and J2 at 1x MIC did not differ significantly; both strains regrew after 4 hours. CZA time-kills of the strains at 4x MIC did not significantly differ from those at 1x MIC. Data are presented as mean log_10_(CFU/mL) ± standard error from four independent experiments. MVB MICs against both strains J1 and J2 were 0.06 µg/mL. Respective CZA MICs were 1 and 2 µg/mL.

#### Capsular polysaccharide (CPS) and mucoviscosity

To investigate CPS content, we quantified uronic acid concentrations in representative strains from patients A (A1, A4, A8), G (G1, G7) and J (J1, J2, J5, J6, J7) (Table 4). As expected, strains J2, J5, J6 and J7 had significantly lower CPS content than strain J1 (mean uronic acid: 33.3 *vs.* 76.2 nmole/mL; p=0.004) or strains from patients A and G (mean uronic acid: 144.2 and 99.4 nmole/mL, respectively; p=0.009 and 0.02). We next evaluated mucovicosity by measuring supernatant turbidity (optical density at 600 nm (OD_600_)) after low-speed centrifugation. Strains J2, J3, J6 and J7 exhibited significantly less mucoviscosity than strain J1 (mean OD_600_: 0.44 *vs.* 0.69; p=0.004) or strains from patient A and G (means: 0.70 and 0.69, respectively; p<0.0001 and p=0.0001). Differences in mucoviscosity were also clearly visualized within tubes following centrifugation (Figure 6). There were no significant within-host differences among A strains or G strains in either uronic acid concentrations or supernatant turbidity.

**Figure 6.**
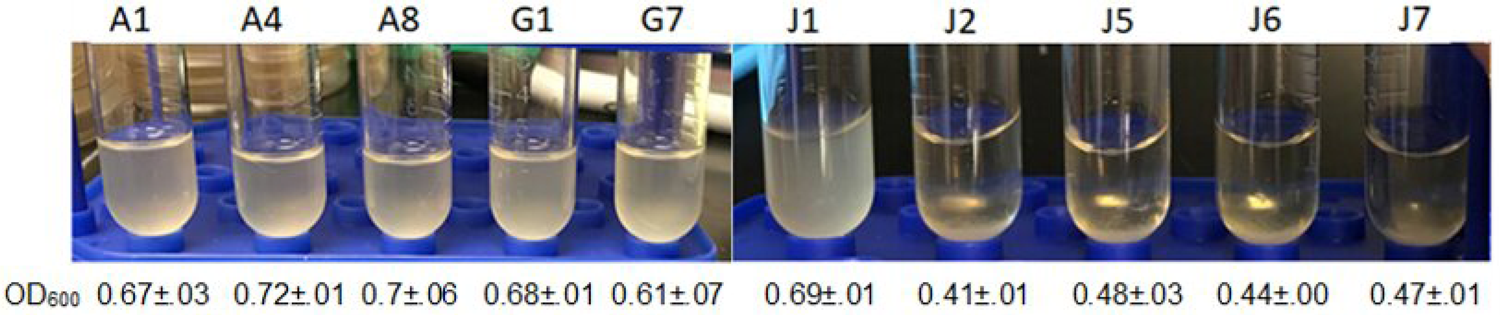
Mucoviscosity of carbapenem-resistant *Klebsiella pneumoniae* strains from three patients (A, G, J). Mucoviscosity was measured as optical density at 600 nm (OD_600_) of supernatants following low-speed centrifugation of strains in Luria-Bertani liquid medium. Photos of representative post-centrifugation tubes for respective strains are shown. OD_600_ data (mean ± standard error of mean) for each strain appear below the photos. Strains with less mucoviscosity are less turbid in tubes and have lower OD_600_ than strains with higher mucoviscosity. Strain J2, J5, J6 and J7 exhibited significantly less mucoviscosity than J1, or strains from patients A and G.

**Table 4.** Capsular genotypes and *in vitro* phenotypes of carbapenem-resistant *Klebsiella pneumoniae* strains from three patients (A, G, J).

| Phenotype | Strains from patient A |  |  |  | Strains from patient G |  |  | Strains from patient J |  |  |  |  |  |
| --- | --- | --- | --- | --- | --- | --- | --- | --- | --- | --- | --- | --- | --- |
|  | A1 | A4 | A8 | p-values | G1 | G7 | p-values | J1 | J2 | J5 | J6 | J7 | p-values |
| Capsule synthesis locus | KL107 | KL107 | KL107 | NA | KL107 | KL107 |  | KL107-2.2Kb | KL107-2.2-11Kb | KL107-2.2-3.6Kb | KL107-2.2-6Kb | KL107-2.2-2.7Kb | NA |
| CPS (uronic acid, nmole/mL) | 140.8±4.8 | 159.6±10.9 | 134.6±15.4 | NS (0.07) | 109.2±4.1 | 114.4±7.8 | NS (0.10) | 76.2±5.8 | 28.4±3.0 | 39.3±5.9 | 29.6±3.8 | 35.9±1.3 | 0.0001 |
| Mucoviscosity (OD<sub>600</sub>, mean ± S.D.) | 0.67±0.03 | 0.72±0.01 | 0.68±0.04 | NS (0.20) | 0.68±0.01 | 0.61±0.03 | NS (0.10) | 0.69±0.01 | 0.41±0.001 | 0.48±0.03 | 0.44±0.01 | 0.47±0.01 | <0.0001 |
| % serum kill (mean ± SD) | 87.0±3.4 | 52.2±1.27 | 86.6±0.8 | NS (p=0.07) | 56.2±4.4 | 52.9±4.5 | NS (p=0.86) | 85.7±2.3 | 100%±0 | 100±0 | 100±0 | 100±0 | 0.01 |
| % macrophage kill (mean ± SD)* | 40.0±4.3 | 35.6±7.3 | 36.8±4.8 | NS (p=0.47) | 37.6±4.9 | 22.7±9.7 | NS (p=0.23) | 55.4±8.6 | 52.8±8.4 | 58.8±9.0 | 56.3±10.9 | 52.1±9.2 | NS (p=0.52) |

Data are presented as mean values from at least 3 independent experiments.
CPS; capsular polysaccharide; nmol/mL: nanomole/milliliter; OD; optical density; SD: standard deviation; NA: not applicable; NS: non-significant; kb: kilobases
\*RAW264.7 macrophages

#### Serum and macrophage killing

CPS has been shown to protect bacteria from complement-mediated killing in serum (34). Strains J2, J5, J6 and J7 were completely killed upon serum incubation; these strains were significantly more susceptible to serum killing than strain J1 or strains from patients A (A1, A4, A8) or G (G1, G7) (all p≤0.01) (Table 4). There were no significant differences in susceptibility to serum killing among A strains or G strains. There were no significant within-host strain differences in susceptibility to macrophage killing.

#### Virulence during bloodstream infections of mice

To examine virulence *in vivo*, we infected cyclophosphamide- and cortisone-treated mice intravenously with strains from patients A (A1, A4, A8), G (G1, G7) or J (J1, J2, J5). Outcomes, measured as mortality or tissue burdens within spleen, kidney and liver, were worse for mice infected with strains A1 and A4 than A8 (Figure 7). There were no differences in outcomes of mice infected with G1 or G7. Mice infected with strain J1 had worse outcomes than mice infected with J2 or J5, by both mortality and tissue burden endpoints.

**Figure 7.**
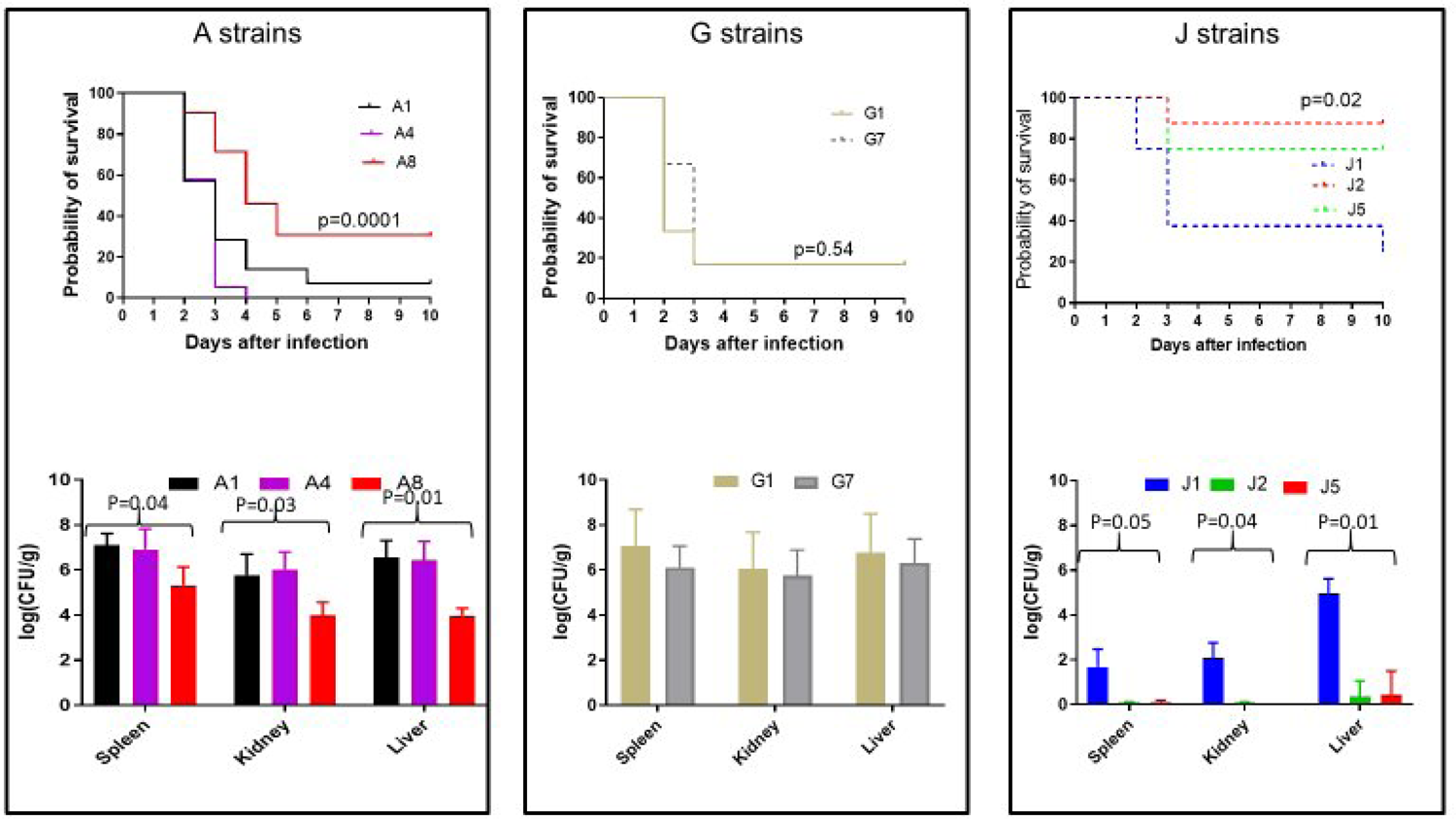
Mortality and tissue burdens of immunosuppressed mice during hematogenously disseminated *Klebsiella pneumoniae* infections. Cyclophosphamide- and cortisone-treated mice were infected intravenously with strains indicated (mortality studies: 1×10^5^ colony forming units (CFUs) per strain; tissue burden studies: 1×10^4^ CFUs per strain). Mortality and tissue burden data are presented in top and bottom panels, respectively. Tissue burdens (log_10_CFU/g tissue, presented as mean ± standard error of mean) were determined in spleens, kidneys and livers at 48 hours. Differences in mortality were compared using Mantel-Cox log rank test. Differences in tissue burdens were compared using Kruskal-Wallis (for A and J strains; comparisons between 3 strains) or Mann-Whitney tests (for G strains; comparisons between two strains). There were significant differences in outcomes (mortality, tissue burdens) among mice infected with strains from patients A and J. There was no difference in outcomes among mice infected with strains from patient G.

To corroborate that deletion of *wzc*, as observed in J2 and several other J strains, contributed to attenuated virulence, we used CRISPR-cas9 to create an isogenic gene disruption strain in the background of A1 and infected immunosuppressed mice intravenously. We used strain A1 rather than J1 to create the mutant because the latter contains other capsular mutations that might confound results. *wzc* mutant and paired A1 parent strains exhibited similar growth rates *in vitro* (data not shown). The null mutant had significantly reduced CPS content (data not shown), and it caused significantly lower burdens within spleens, kidneys and livers of mice during BSIs compared to those caused by the parent strain (Supplemental Figure 3).

## DISCUSSION

This study is the first genome-wide analysis of within-host diversity of *K. pneumoniae* recovered from individual patients with BSIs. We showed that positive blood cultures were comprised of genetically heterogeneous ST258 *K. pneumoniae* populations, with strains differing by core genome SNPs, presence or absence of specific genes (including those involved in antibiotic resistance, capsular synthesis and other processes relevant to pathogenesis), and/or plasmid content. Moreover, we demonstrated that genetically diverse strains exhibited unique phenotypes that are potentially important during BSIs, including differences in antibiotic responses, CPS and mucoviscosity, resistance to serum killing, and ability to cause organ infections or mortality *in vivo*. This diversity was not appreciated by standard clinical microbiology laboratory approaches, in which a single strain was selected from positive blood cultures for further characterization. Rather, diversity was unmasked only by studying strains from multiple, morphologically indistinguishable colonies. Our data support a new, population-based paradigm for CRKP BSIs.

Genetically diverse ST258 *K. pneumoniae* were recovered from first positive blood cultures in all 6 patients we investigated. To our knowledge, only one other study has assessed bacterial genetic diversity during BSIs by whole genome sequencing of strains from multiple, morphologically indistinguishable colonies. In that study, *Staphylococcus aureus* genetic variants of the same ST were identified in 36% of bacteremic patients (12). Investigators did not report whether genetic variant strains exhibited distinct phenotypes. The higher prevalence of genetic diversity we observed may reflect sequencing strains from 10 rather than 3-5 colonies, differences in comparative genomic analytic methods, and/or particularly strong selection pressures encountered by patients with CRE infections. Genetic variants are increasingly recognized among bacterial strains during colonization and chronic infections of non-sterile sites, including the GI tract (*H. pylori*, *Enterococcus* spp.) (16, 17, 19, 40), lungs (*Pseudomonas aeruginosa*, *Burkholderia* spp., *Mycobacterium* spp.) (14, 18, 20), nasal cavity (*S. aureus*) (12, 13, 22), and skin (*Staphylococcus epidermidis*) (41). Emergence of diversity might be expected during such long-term interactions with the host (19, 21, 40, 42), but it is more surprising in acute infections of a normally sterile site. In one study, capsular gene mutant subpopulations were identified in 10% of *K. pneumoniae*-positive urine cultures, including samples associated with acute urinary tract infections (UTIs) in 2 patients (37). In fact, diversity may have been underestimated in these cultures since investigators screened for hypermucoid phenotypes, rather than employing a sequence-first approach.

Pathogens face unique selection pressures in the bloodstream compared to those encountered as GI commensals or at other sites of colonization. These pressures may select for outgrowth of variant strains within the population that are better able to persist and proliferate as opportunistic pathogens (40, 42). The capsule is the major virulence factor in *K. pneumoniae* and other Enterobacterales (34). Capsular gene mutations were identified in strains from 4 of 6 patients (A, D, F, J), each of which had a KL107 capsule type. In 3 patients, there was within-host diversity of capsular mutant strains, including variants with non-synonymous *wzc* SNPs (A, J) and various gene deletions (D, J). J strains with extensive capsular gene deletions were significantly attenuated in CPS content and mucoviscosity, more susceptible to serum killing, and less virulent during hematogenously disseminated infections of mice than strain J1, which had more limited capsular deletion. Using isogenic ST258 strains, we showed that disruption of *wzc*, as in several J strains, led to significantly reduced CPS content and lower tissue burdens in mouse infections. Capsular gene mutant *K. pneumoniae* of various STs that exhibit hypermucoid or hypomucoid phenotypes have emerged repeatedly and independently in clinical cultures; 10% of ST258, clade 2 *K. pneumoniae* genomes in the National Center for Biotechnology Information RefSeq database carry non-synonymous mutations in *wzc* or *whaP* capsule genes (37). Our findings of attenuated virulence for hypomucoid strains with *wzc* disruption or more extensive capsular mutations are broadly consistent with prior observations that hypermucoid *K. pneumoniae* caused greater lethality in mouse BSIs (37).

Patient J was treated with meropenem and other antibiotics prior to diagnosis of CRKP BSI. We showed that strain J2, which had the largest capsular gene deletion among J strains, re-grew after 4-24 hours of exposure to meropenem-vaborbactam or ceftazidime-avibactam *in vitro*, despite MICs in the susceptible range. In contrast, the drugs exerted prolonged bactericidal activity against strain J1. These findings are consistent with those from a previous study in which a capsule-deficient *K. pneumoniae* strain was more tolerant than a wild-type strain to meropenem-vaborbactam within bladder epithelial cells *in vitro* (37). It is plausible that capsular mutations that reduce intrinsic *K. pneumoniae* virulence during BSIs may afford advantages during antibiotic treatment. The data carry potential clinical importance, since meropenem-vaborbactam or ceftazidime-avibactam are drugs of choice against KPC-producing CRKP infections (38). Of note, the meropenem-vaborbactam-tolerant, capsule-deficient *K. pneumoniae* strain from the earlier study demonstrated enhanced virulence in untreated mice with chronic UTIs (37). Therefore, capsular mutations that reduce fitness in some environments *in vivo* may be advantageous in other environments, independent of contributions to antibiotic responses. Taken together, data in the present and past studies attest to the complex, multi-factorial nature of *K. pneumoniae* virulence. Along these lines, strain A4 lacked numerous virulence genes found in other A strains, including those encoding aerobactin and siderophores, but it nevertheless caused higher tissue burdens and greater mortality during hematogenously disseminated infections of mice.

The clinical significance of CRKP diversity shown here is unknown, and merits further investigation. A strength of our study design is that CRKP strains were collected as blood cultures were being processed according to standard clinical microbiology laboratory practices. The possible impact of these practices is highlighted in patient G, who would have been diagnosed with BSI due to carbapenem-susceptible *K. pneumoniae* rather than CRKP if G7 was randomly selected as index strain instead of G1. Indeed, studies of patients diagnosed with more susceptible *K. pneumoniae* BSIs than were identified in our patients would allow investigators to address how often clinical laboratories fail to identify resistant variants (i.e., heteroresistance) within the population, and whether such events lead to treatment failures (43). Future longitudinal studies should assess treatment responses, patient outcomes, and endpoints such as emergence of *de novo* antibiotic resistance, selection for pre-existing resistant strains, or adaptive bacterial evolution. If bacterial diversity is proven to be clinically relevant, microbiology laboratory practices will need to be modified. At present, our findings and those of studies showing bacterial genetic diversity at non-bloodstream sites have important implications for molecular epidemiologic investigations of infectious outbreaks and nosocomial transmission of pathogens. Data here and in our previous study of clinical and environmental Mucorales suggest caution in relying upon core genome SNP phylogeny as the sole tool in defining differences between strains (44). In both studies, comprehensive pan-genome analyses revealed variations that were not apparent with core genome comparisons.

It is unclear whether within-host diversity we describe stemmed from one-time inoculation of a mixed population from the GI tract or other portal, serial introduction of different strains, or evolution within the bloodstream or blood cultures. For several reasons, we believe it is most likely that diversity was generated within the GI tract. Genetic and phenotypic diversity is well-described within GI-colonizing populations of other enteric bacteria (17, 19, 40, 42). Most CRKP BSIs are caused by GI-colonizing strains, and patients at-risk for CRE infections typically encounter intense and long-term selection pressures for microbial diversification, including repeated exposure to broad-spectrum antibiotics (6–8, 45). Our detection of mutations in biologically plausible targets that were previously described among *K. pneumoniae* clinical isolates recovered from diverse body sites, such as antibiotic resistance, capsular biosynthetic and porin genes, support the validity of our findings and suggest they reflect diversity *in vivo* (6, 35, 37, 45). In the future, metagenomic sequencing may afford in-depth coverage of microbial variants in samples directly from sites of infection. Currently, however, metagenomic sequencing of bacteria within blood is limited by overwhelming predominance of host DNA and low concentrations of microbial DNA, need for target amplification, and challenges in assigning sequence variations to individual strains (46). Follow-up studies are warranted to compare CRKP diversity at GI and other sites of colonization with that observed during BSIs. We acknowledge that our findings cannot necessarily be extrapolated to other *K. pneumoniae* STs or bacterial species. In ongoing studies, we have demonstrated genetic diversity among non-ST258 CRKP and carbapenem-resistant *K. michiganensis* from blood cultures (unpublished data) (47).

In conclusion, we identified genotypic and phenotypic variant strains of ST258 *K. pneumoniae* from blood cultures of individual patients. Clinical implications of such genetic and phenotypic diversity during BSIs and other infections will be defined in upcoming years, with potentially profound implications for medical, clinical microbiology laboratory and infection prevention practices, and for better understanding of emergence of antibiotic resistance and pathogenesis.

## MATERIALS AND METHODS

### Strains and growth conditions

Six patients with CRKP BSI were identified at the University of Pittsburgh Medical Center between April 2017 and August 2018. First positive blood culture bottles from each patient were obtained from the clinical laboratory immediately after routine microbiological work-up. We streaked 25 µL of broth from culture bottles onto blood agar plates (5% Sheep Blood in Tryptic Soy Agar), and randomly picked 9 morphologically indistinguishable single colonies. Each colony was subcultured onto Mueller-Hinton (MH) agar plates. Following overnight growth at 37 °C, a single strain per original colony underwent DNA extraction (PureLink™ Genomic DNA Mini Kit; Fisher); remaining colonies and confluent growth were frozen in 20% glycerol at −80 °C. We also obtained the index strain from each patient that was isolated by the clinical laboratory, and extracted DNA and made frozen stock using the methods above. “Strain” is this study is defined as a CRKP isolate from a single colony that was shown to be genetically distinct by comprehensive WGS analyses.

### Short-read WGS and analyses

Sixty strains (10 per patient) were sequenced using Illumina HiSeq. Raw short-reads were quality-trimmed and *de novo* assembled into contigs using Shovill v1.1.0 (https://github.com/tseemann/shovill). We evaluated genome assembly quality by Quast v5.0.2. Draft genomes were screened for contamination and annotated using MASH v2.3 and Prokka v1.14.5, respectively. Species identification, and K & O and sequence typing were performed using Kleborate v2.0.4. Antibiotic resistance, virulence and plasmid replicon type genes were detected in assembled genomes by ABRicate v1.1.0, using National Center for Biotechnology Information (NCBI), virulence factor (VFDB, http://www.mgc.ac.cn/VFs/) and dtu.dk/services/PlasmidFinder/). Variant calling and core-genome SNP alignment were produced by Snippy v4.6.0 (https://github.com/tseemann/snippy), using *K. pneumoniae* 30660/NJST258_1 (GenBank assembly accession: GCA_000598005.1) as reference. We used the alignment of core SNPs as input for MEGA11 to generate a maximum likelihood phylogenetic tree. The phylogenetic tree and associated metadata were visualized using iTOL v5.6.3. A pan-genome was constructed by Roary v3.13.0 using default settings, paralog splitting on, and 95% minimum identity for BLASTp. Core genes were defined as those present in 99% of strains, which prevented the outgroup strain from affecting core genome estimation.

### Long-read WGS and hybrid assemblies

Strains underwent long-read Oxford Nanopore sequencing using MinION. In brief, DNA was isolated from overnight cultures using a MasterPure™ Gram Positive DNA Purification Kit (Epicentre, USA). Nanopore libraries were prepared using Ligation Sequencing Kit (SQK-RBK109) and sequenced by an R9.4 flow cell using a MinION MK1B device. We used Unicycler to hybrid assemble MinION and Illumina reads. Hybrid assemblies were visualized using Bandage v0.9.0 (48). Variant calling was performed using Snippy v4.6.0.

### Antibiotic susceptibility testing and time-kills

MICs were determined in duplicate by the Clinical and Laboratory Standards Association reference broth microdilution method. *K. pneumoniae* ATCC700603 served as internal quality control. Time-kill assays (4 replicates) were performed as previously described, using a single bacterial colony grown overnight in 4 mL MH Broth (MHB) (49). Samples were taken at 0, 4, 8, 12 and 24 hours, serially diluted, spread on plates, and incubated at 37°C. Colonies were counted after incubation at 37 °C for 24 hours.

### *ompK36* reverse transcription-polymerase chain reaction (RT-PCR) and sodium dodecyl sulfate–polyacrylamide gel electrophoresis (SDS-PAGE)

RT-PCR and SDS-PAGE were performed as previously described (49). Relative quantities of mRNA from each gene were determined by comparative threshold *C_T._* Expression of *ompK36* was normalized to that of *rec*A and *rpo*D. Outer membrane proteins were analyzed in 12% SDS-PAGE and strained with silver stain kit (Bio-Rad). These experiments were performed in triplicate, as were each of *in vitro* phenotypic assays below.

### CPS quantitation

CPS was extracted and uronic acid concentrations were quantified (nMole/10^9^ CFU) using established methods (50).

### Mucoviscosity

Mucoviscosity was assessed by low-speed centrifugation of CRKP strains grown overnight in Luria-Bertani liquid medium at 37 °C. Cultures were normalized to optical density at 600 nm (OD_600_)=1 and centrifuged at 100x*g* for 20 minutes at 22 °C (Marathon 3000R, swinging bucket, Fisher Scientific). OD_600_ of supernatant was determined (BioMate 3 Thermo Spectronic, Fisher Scientific).

#### Serum killing (51)

Overnight cultures were diluted 1:100 and grown to mid-exponential phase. An inoculum of 2.5×10^4^ CFUs of bacteria in 25 uL was mixed with 75 μL human serum from healthy volunteers (Cat# BP2657100; Fisher Scientific). The mixture was incubated at 37 °C and aliquots were taken at baseline and 1h to calculate viable CFUs. Average survival percentage was plotted against time.

#### Macrophage killing (52)

*In vitro* killing of strains was investigated using the RAW264.7 macrophage cell line. In a 96-well plate, 8×10^4^ macrophages were re-suspended in DMEM, seeded in 3 wells and incubated overnight at 37°C. After 12 hours, 1.6×10^6^ CRKP CFUs were added onto the monolayer at time 0, and incubated for 15 min at 37 °C. Bacteria were washed 3 times with DMEM and macrophages were lysed with H_2_O. The number of intracellular bacteria was determined by serial dilutions. Intracellular killing was based on the decrease of viable bacteria 30 minutes after initial co-incubation.

### Mouse infections

Male ICR CD1 mice weighing 20–25 g (Harlan) were immunosuppressed with 2 doses of intraperitoneal cyclophosphamide (150 mg/kg 4 days prior to infection, 100 mg/kg 1 day prior to infection) and 2 doses of subcutaneous cortisone (20 mg/kg 4 days and 1 day prior to infection). For mortality studies, mice (12/group) were infected via lateral tail vein with 1×10^5^ CRKP suspended in 100 µL of saline. Mice were followed until they were moribund, at which point they were sacrificed, or for 10 days. Survival curves were calculated according to the Kaplan-Meier method using the PRISM program (GraphPad Software) and compared using Mantel-Cox log-rank test. For tissue burdens, 1×10^4^ CRKP were injected via lateral tail vein. Mice (12/group) were sacrificed at 48 hr post infection. Differences in tissue burden were presented as mean (±standard error) log_10_ CFU/gram of tissue, and analyzed by Kruskal-Wallis (for studies with three strains; A and J) or Mann-Whitney (for studies with 2 strains; G and isogenic *wzc* mutants) tests. *p*<0.05 was considered statistically significant in all phenotypic studies.

### CRISPR-Cas9 deletion of *wzc*

CRISPR-Cas9-mediated *wzc* deletion was conducted as previously described (53). *wzc* specific guide RNA (gRNA) (GGTTTTGATGTAATACAGAG) was designed and inserted into plasmid vector pSgKp-Rif (54). *wzc* gRNA pSgKp-Rif and a 90-nt synthesized template (AAAGCATGGGCGGAAAAATTAGCAAGTTAATTCAGGAAAATATACAGAAGTGTTTTCAACAAAAGCCGGATTTAGATAAATATTTAGATT) were electroporated into pCasKp-harboring CRKP A1 competent cells to create the knockout strain.

## DATA AVAILABILITY

Raw reads for 60 ST258 *K. pneumoniae* strains were submitted to the Sequence Read Archive (SRA). BioProject: PRJNA826066. BioSample numbers: SAMN27547637-SAMN27547696.

## ACKNOWLEDGMENTS

This project was supported by NIH grants R21AI160098 (MHN), R21 AI152018 (CJC) and R01AI090155 (BNK).

## AUTHORS’ CONTRIBUTIONS

SJC: Extracted strain DNA, conducted RT-PCR, SDS-PAGE, other *in vitro* and mouse experiments, interpreted data, drafted manuscript, and formatted tables and figures. GF: Whole genome sequence data analyses, assisted with drafting and editing manuscript, and prepared genomics tables and figures. LC: Whole genome sequencing and accompanying data analyses, performed CRISPR-cas9 gene disruption, and assisted with editing manuscript. GL, BH, AN, ED, KMS: Conducted experiments in conjunction with SJC. RKS: Blood culture collection and assisted with editing manuscript. TC: Conducted CRISPR-cas9 gene disruption in conjunction with LC. BNK: Assisted with whole genome sequence analyses and editing manuscript. MHN, CJC: Study conception and design, oversaw experiments and data analyses, and re-drafted and edited manuscript.

## SUPPLMENTAL FIGURE LEGENDS

**Supplemental Figure 1.** *ompk36* sequences in carbapenem-resistant *Klebsiella pneumoniae* strains from patient G (G3, G4, G6). Strains G4 and G6 carried mutations in the promoter region of *ompk36*. *ompk36* sequence in strain G3 was similar to that of the remaining G strains. rbs: ribosome binding site

**Supplemental Figure 2.** *ompk36* expression and ompK36 porin production by carbapenem-resistant *Klebsiella pneumoniae* strains from patient G (G1, G4, G6, G7). Relative gene expression is shown on the left for each strain in presence and absence of meropenem-vaborbactam (MV), as determined by reverse transcription-polymerase chain reaction. Protein production in absence of drug exposure is shown on the right, as determined by 12% sodium dodecyl sulfate–polyacrylamide gel electrophoresis. The lane marked “Ladder” contains size markers in kiloDaltons (kDa).

**Supplemental Figure 3.** Tissue burdens of mice infected intravenously with a carbapenem-resistant *Klebsiella pneumoniae wzc* null mutant strain. Mice were infected intravenously with a parent (pCasKp-harboring A1 competent cells; black bars) or *wzc* isogenic null mutant strain created in the parent background by a CRISPR-cas9 method (checkerboard bars). Tissue burdens in respective organs (log_10_ CFU/gm ± standard error) at 48 hours post-infection appear above bar graphs. Differences in tissue burden were analyzed using Mann-Whitney test; *p*<0.05 was considered statistically significant. Disruption of *wzc* resulted in significant reductions in *K. pneumoniae* burdens within spleens, kidneys and livers.

## SUPPLMENTAL TABLE LEGENDS

**Supplemental Table 1.** Assembly data for carbapenem-resistant *Klebsiella pneumoniae* strains.

**Supplemental Table 2.** Pan-genome matrices for carbapenem-resistant *Klebsiella pneumoniae* strains. Please refer to Excel file in supplemental materials.

**Supplemental Table 3.** Antibiotic minimum inhibitory concentrations against carbapenem-resistant *Klebsiella pneumoniae* strains from three patients (A, G, J). Susceptibility testing was performed using the Clinical and Laboratory Standards Institute reference broth microdilution method. Bolded MICs within shaded boxes differ by ≥4-fold from MICs against other strains. MIC: minimum inhibitory concentration; MEM: meropenem; MVB: meropenem-vaborbactam; CAZ: ceftazidime; CZA: ceftazidime-avibactam; TET: tetracycline; GEN: Gentamycin

